# Terpenoid balance in *Aspergillus nidulans* unveiled by heterologous squalene synthase expression

**DOI:** 10.1101/2023.10.20.563295

**Authors:** Sung Chul Park, Breanne N. Steffan, Fang Yun Lim, Raveena Gupta, Fatma Ayaloglu Butun, Hongyu Chen, Rosa Ye, Timothy Decker, Chengcang C. Wu, Neil L. Kelleher, Jin Woo Bok, Nancy P. Keller

**Author notes:** Corresponding authors. Email: Jin Woo Bok, Nancy P. Keller.

## Abstract

Filamentous fungi produce numerous uncharacterized natural products (NPs) that are often challenging to characterize due to cryptic expression in laboratory conditions. Previously, we have successfully isolated novel NPs by expressing fungal artificial chromosomes (FACs) from a variety of fungal species into *Aspergillus nidulans*. Here, we demonstrate a new twist to FAC utility wherein heterologous expression of a *Pseudogymnoascus destructans* FAC in *A. nidulans* altered endogenous terpene biosynthetic pathways. In contrast to wildtype, the FAC transformant produced increased levels of squalene and aspernidine type compounds, including three new nidulenes (**1**–**2, 5**), and lost nearly all ability to synthesize the major *A. nidulans* characteristic terpene, austinol. Deletion of a squalene synthase gene in the FAC restored wildtype chemical profiles. The altered squalene to farnesyl pyrophosphate ratio leading to synthesis of nidulenes and aspernidines at the expense of farnesyl pyrophosphate derived austinols provides unexpected insight into routes of terpene synthesis in fungi.

**Teaser:** Reshaping terpenes: Heterologous FAC expression reroutes terpene pathways.

## Introduction

Fungi produce a wide variety of natural products (NPs, also called secondary metabolites), some of which have important medicinal properties as drug candidates (*1–3*). Among these metabolites, meroterpenoids have emerged as a class of secondary metabolites that possess complex structures and diverse bioactivities (*4, 5*). Meroterpenoids are formed through the mixing of terpene biosynthetic pathways with other biosynthetic origins such as polyketide or non-ribosomal peptide pathways resulting in a high degree of structural diversity (*4, 6*). These compounds have attracted significant attention in drug discovery and development due to promising pharmacological activities, including antifungal, antibacterial, antiviral, anti-inflammatory, and anticancer properties (*7*). More than 1,500 novel fungi-derived meroterpenoids have been isolated and published since 2009 (*8*). As such, understanding the biosynthesis, structure, and biological activities of fungal meroterpenoids is of great interest to the field of drug discovery.

One limitation of current drug discovery in fungi is the difficulty in expressing silent fungal biosynthetic gene clusters (BGCs) under laboratory conditions to produce NPs for drug candidate screening (*9*). Fungal artificial chromosomes (FACs) represent a genetic tool to produce unknown fungal NPs by transforming randomly sheared genomic DNA containing fungal BGCs in heterologous hosts, such as *Aspergillus nidulans* (*10*). FACs offer the advantages of the easy association of mass spectrometry (MS) detectable metabolites with their transformed BGCs (*10*). Many FAC-derived NPs are cryptic metabolites not typically produced by the endogenous species, resulting in an increased potential to discover novel fungal metabolites and related BGCs (*11*). For example, benzomalvins A/D (*11*), valactamide A (*11*), and terreazepine (*12*) and their BGCs were newly identified by applying FACs transformation into *A. nidulans*.

*A. nidulans* is a renowned heterologous expression host due to the ease of genetic manipulation first established in the late 1940s (*13*) and easily adapted to new genetic tools in the subsequent decades (*14*). Further, studies in this fungus in the last 30 years have linked over 30 BGCs to specific NPs (*15*) which simplifies the identification of heterologous NPs such as those expressed from FACs. Yet, *A. nidulans* contains over 70 BGCs (*16*) suggesting that many new compounds still remain to be identified in this fungus. As a twist in the use of FAC technology, here we present a new utility of this technology to induce new endogenous NPs in the expression host. By expressing a *Pseudogymnoascus destructans* squalene synthase (SqsA) containing FAC in *A. nidulans*, we were able to induce biosynthesis of novel endogenous NPs in this host. Instead of producing *P. destructans*-derived metabolites, expression of *sqsA* in *A. nidulans* served as a terpene regulating tool amplifying and redirecting host-derived meroterpenoid pathways. *A. nidulans* contains two known meroterpenoid BGCs encoding the dominant austinol family (*8*) and the silent aspernidine family (*17*) of metabolites. Heterologous expression of *P. destructans sqsA* FAC redirects the *A. nidulans* biosynthetic output from austinol synthesis to aspernidine synthesis concomitant with the production of three new and six known compounds, four possessing immunomodulatory properties (Fig. 1). We suggest leveraging *P. destructans sqsA* FAC as a new tool to increase pools of fungal meroterpenoids and a new method to investigate unidentified fungal meroterpenoids such as the nidulenes A–E (**1**–**5**).

**Fig. 1.**
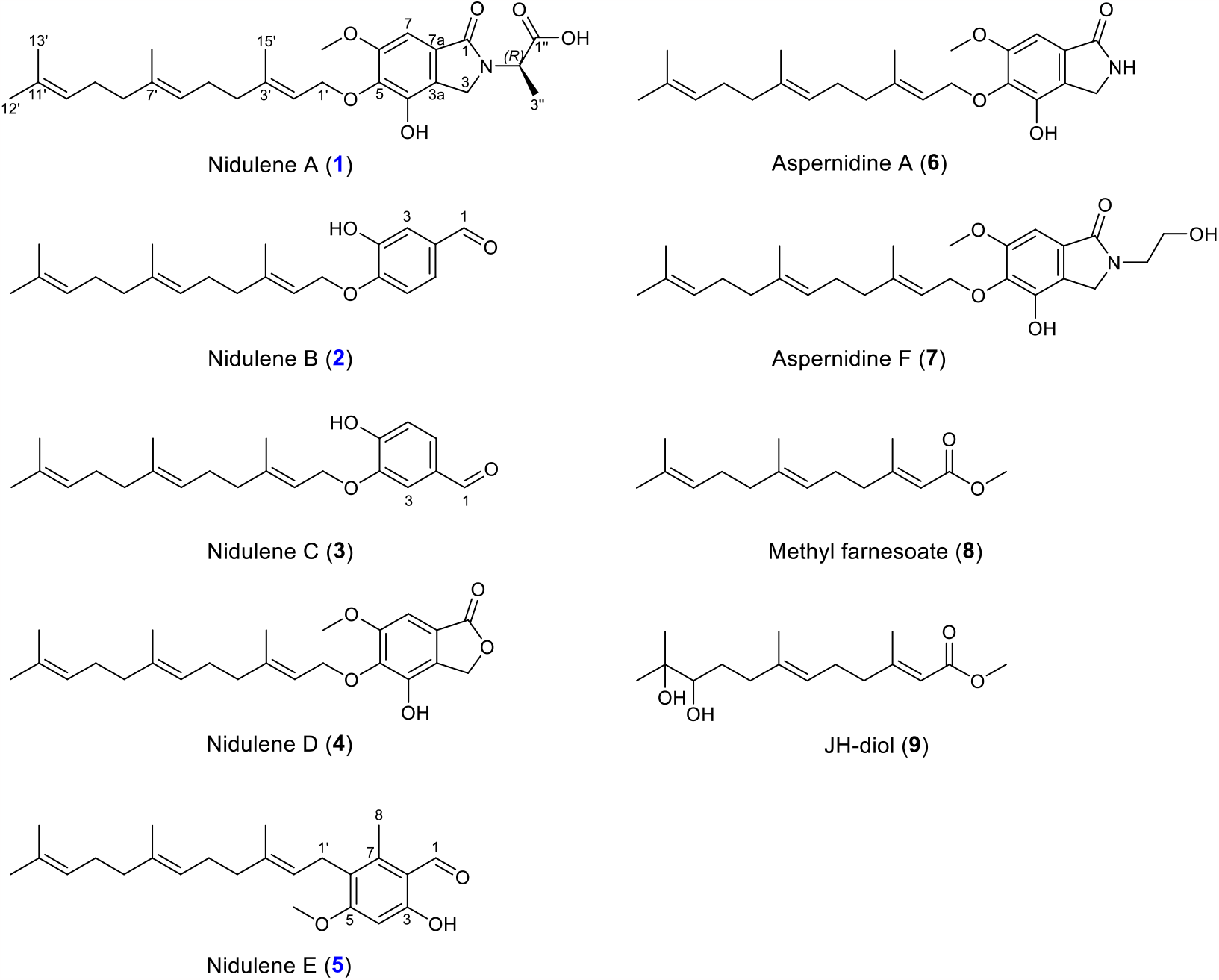
Structure of compounds (1–9). The structure of isolated terpenes from PdFAC1.

## Results

### A *Pseudogymnoascus destructans* FAC induces squalene production in *A. nidulans*

During our search for novel bioactive compounds generated from the FAC transformation of *A. nidulans*, we identified a FAC, PdFAC1, from the fungal bat pathogen *P. destructans* containing the gene VC83_00068 (genebank#: XM_024463765.1) encoding a putative squalene synthase gene, *sqsA*, whose encoded product showed 50% identity to *Saccharomyces cerevisiae* SqsA (*18*) and a 61% identity to the *A. nidulans* SqsA (AN10396). Squalene synthase, also known as farnesyl-diphosphate farnesyltransferase, is a key enzyme in the isoprenoid pathway (*19*) and is required for the synthesis of ergosterol in fungi (*20*) and cholesterol in animals (*21*) (Fig. 2). The primary product of squalene synthase is squalene, which is formed from dimerization of two molecules of farnesyl pyrophosphate.

**Fig. 2.**
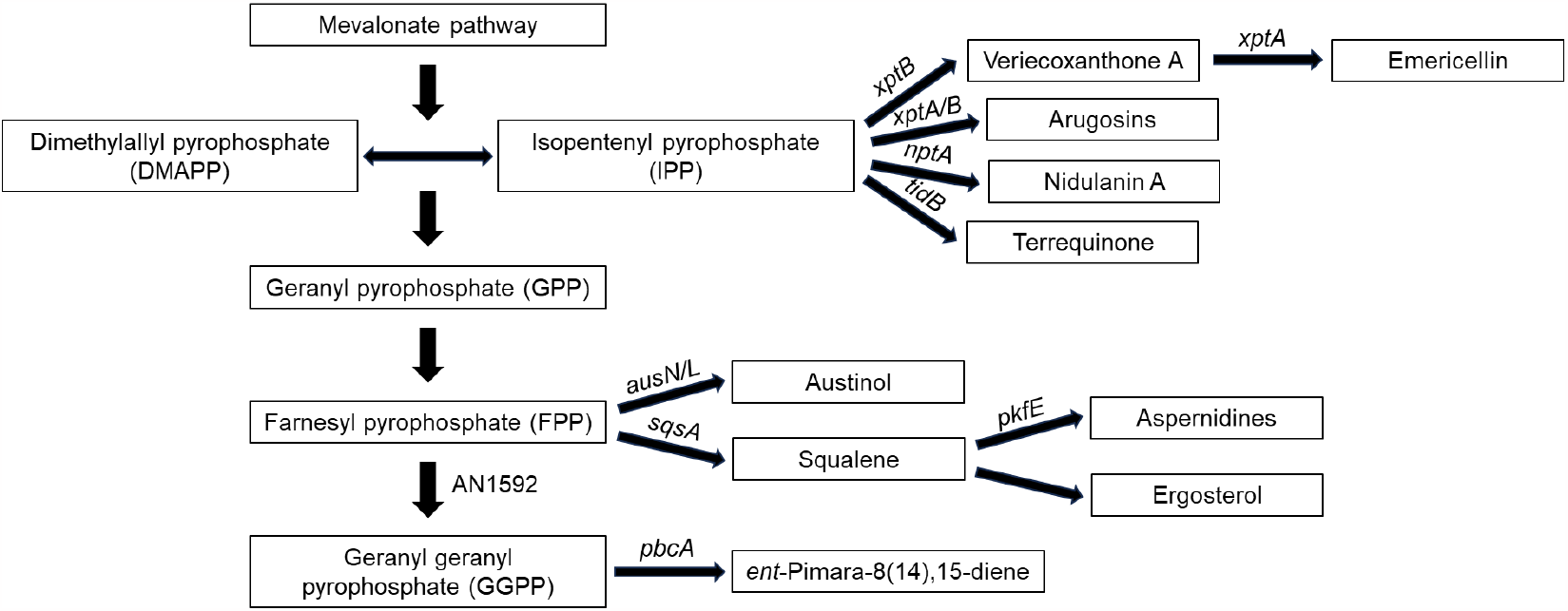
The hypothesized scheme of *A. nidulans* derived terpenes and prenylated metabolites pathways.

To determine if the *sqsA* gene from *P. destructans* affected production of squalene in *A. nidulans*, we compared the squalene production level of a control strain (TJW167), an *A. nidulans* transformant with a PdFAC1 (TJW336), and an *A. nidulans* transformant with PdFAC1 deleted for *sqsA* (TJW337) (Table S1). Metabolite extracts were analyzed by high-resolution UHPLC– MS/MS, and the resulting datasets were compared using the Maven ver.2.0.3 and MZmine ver.3 software, which facilitates comparative analysis of UHPLC–MS/MS data from multiple different strains. The production of squalene was increased ten-fold in TJW336 compared to the control and TJW337, both of which barely produced detectable squalene (Fig. 3). Thus, the squalene synthase gene in PdFAC1 boosted the production level of squalene in *A. nidulans*.

**Fig. 3.**
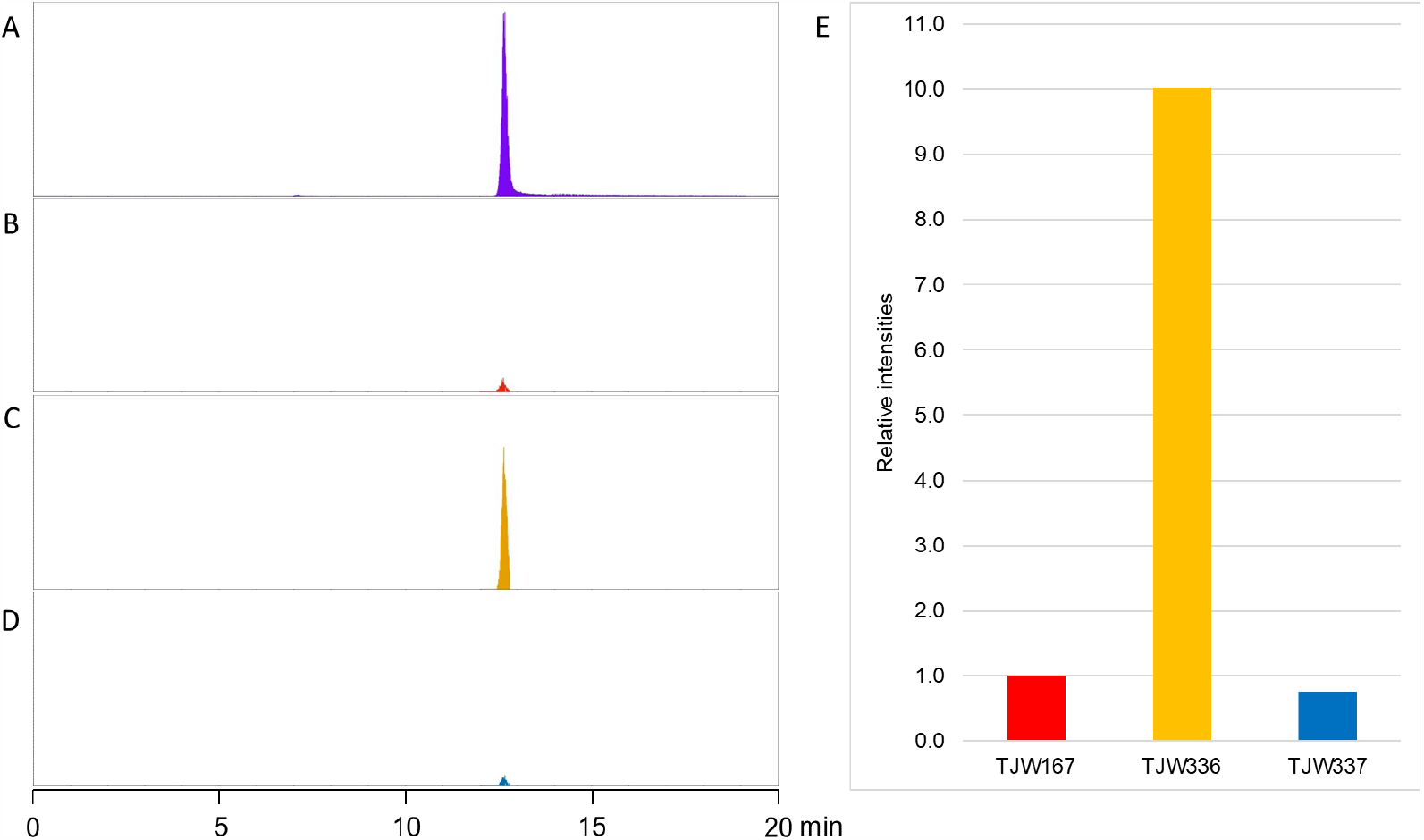
Squalene production is dependent on presence of *P. destructans* derived squalene synthase gene (*sqsA*). **(A)** squalene standard (100 ppm) **(B)** squalene detected from the control strain (TJW167) **(C)** squalene detected from the PdFAC1 transformed *A. nidulans* (TJW336) **(D)** squalene detected from the PdFAC1 transformed *A. nidulans* with deletion of *sqsA* gene (TJW337) **(E)** relative intensities of squalene production from each strain.

### PdFAC1 redirects host meroterpenoid production

Although the role of squalene as an ergosterol precursor is well known (*22*), recent papers have shown that alternative functions of SqsA and SqsA-like enzymes and subsequent squalene synthesis are involved in induction of terpenoids in plants (*23, 24*) and bacteria (*25*). We thus hypothesized that *A. nidulans* transformants expressing PdFAC1 could be altered in terpenoid production. To investigate the effect of SqsA on the chemical composition of host strains, a full range of UHPLC– MS/MS spectra of each strain (control TJW167, TJW336 PdFAC1, and TJW337 PdFAC1Δ*sqsA*) were compared (Fig. 4). While the control strain produced austinol, a cyclized sesquiterpene, as a major *A. nidulans* terpene product, the TJW336 instead produced aspernidine-type meroterpenoids with a linear sesquiterpene attached to the aromatic ring (Fig. 4). From the *sqsA* deleted strain (TJW337), austinol was produced again while the production of aspernidine-type meroterpenoids was gone. Normally the wildtype *A. nidulans* strain used in this study does not produce aspernidines and its BGC was only discovered by screening a protein kinase mutation library where one mutant (Δ*mpk*) produced aspernidines allowing for BGC discovery (*17*). Interestingly both PdFAC1 and Δ*mpk* are greatly diminished in ability to produce austinol while overproducing aspernidines, suggesting a possible shift in the precursor shunt pathways common to both strains. Previous studies have indicated that austinol is derived from farnesyl pyrophosphate (FPP) and 3,5-dimethyl orsellic acid (*17*). Here, it appears that SqsA activity directs FPP into high amounts of squalene in TJW336. We hypothesize that this redirection reduces available farnesyl pyrophosphate pools for austinol synthesis and instead increases squalene pools utilized in synthesis of aspernidine-like metabolites (Fig. 4). Although it has been speculated that the aspernidine terpene tail is derived from farnesyl pyrophosphate (*26*), our data suggests that squalene could be the source of the farnesyl tail, possibly through degradation through oxygenase activity reminiscent to degradation pathways reported in bacteria (*27, 28*).

**Fig.4.**
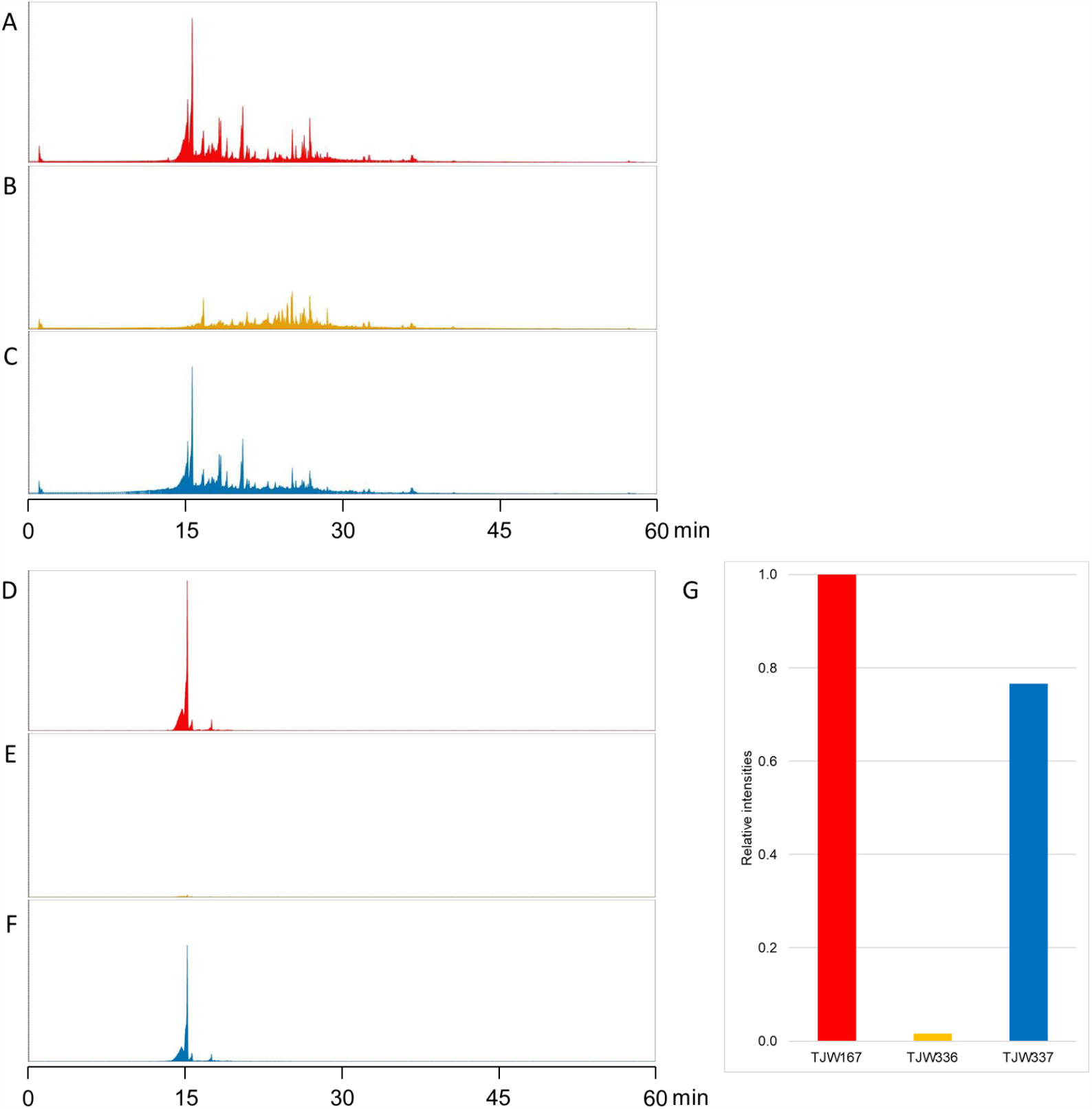
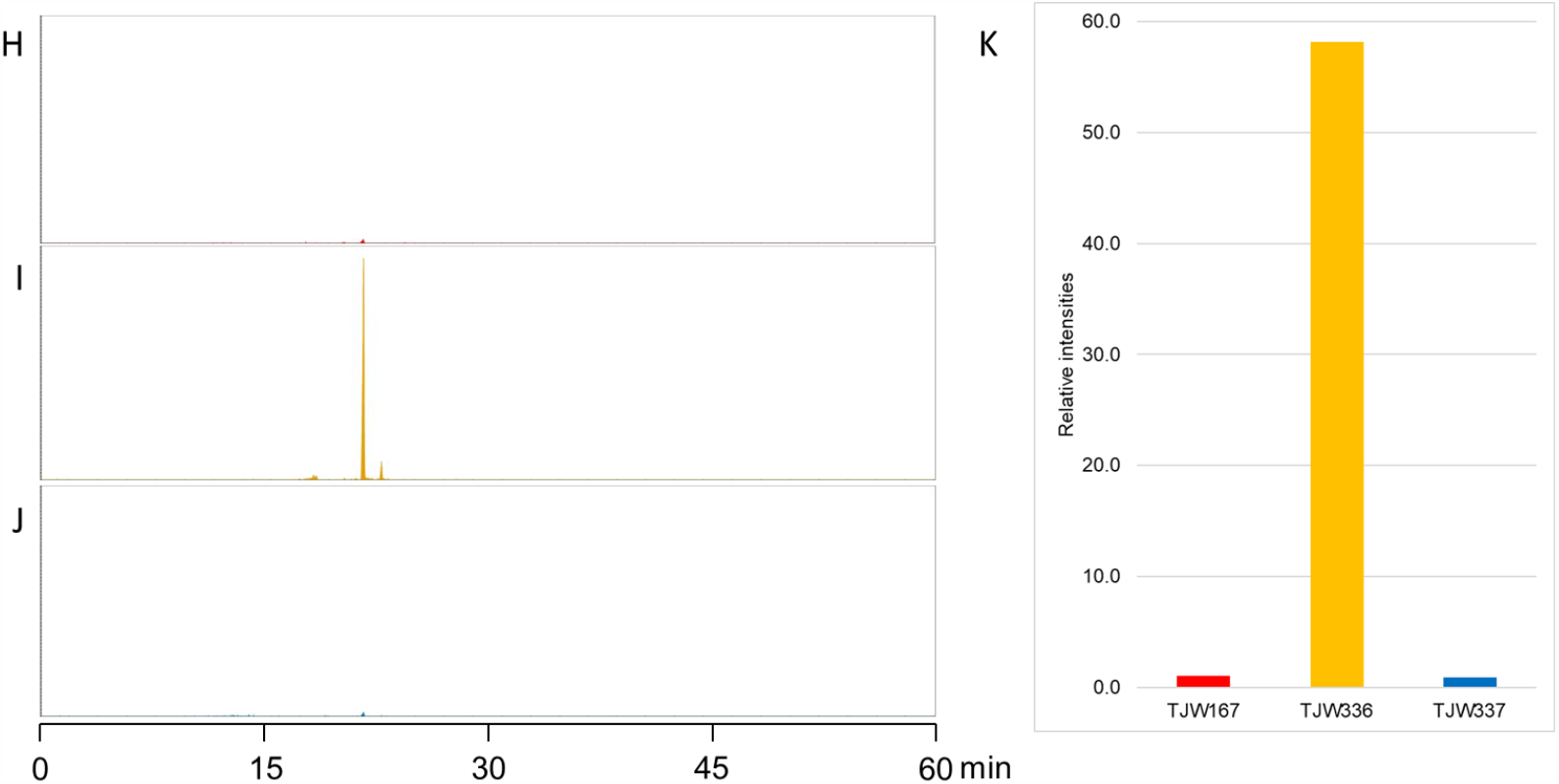
Terpene production is dependent on presence of *P. destructans* derived squalene synthase gene (*sqsA*). **(A–C)** Full scan spectra of TJW167 (red), TJW336 (yellow), and TJW337 (blue), respectively. **(D–G)** Production of austinol decreased to 2% compared to the control strain and recovered after the deletion of *sqsA*. Austinol ([M+H]^+^ = 459.2006) was detected at 15.74 min. **(H–K)** Production of aspernidine A increased to 58-fold compared to the control strain and dropped to the same level with the control strain after the deletion of *sqsA*. Aspernidine A ([M+H]^+^ = 400.2482) was detected at 22.42 min.

### Structure determination of prenylated compounds isolated from TJW336

To determine the structural analysis of the many terpenoids produced in TJW336, each compound was isolated with HPLC. The molecular formula of compound **1** was deduced to be C_27_H_37_NO_6_ with 10 degrees of unsaturation by HR-ESI-MS analysis. The ^13^C NMR and DEPT-135 data of this compound displayed 27 C-atoms, which were assigned to two carbonyl carbons (δ_C_ 173.8 and 167.3), twelve sp^2^ carbons (δ_C_ 153.8–120.8), six methyl carbons (δ_C_ 55.8–15.7), and seven aliphatic carbons (Table 1). Among the seven aliphatic carbons, four carbons were assigned to methylene carbons (δ_C_ 38.9, 38.8, 26.2, and 25.9). The carbon and proton chemical shifts indicated the presence of an oxymethylene (δ_C_/δ_H_ 68.2/4.46), a nitrogenous methylene (δ_C_/δ_H_ 44.5/4.59 and 4.17), and a sp^3^ methine (δ_C_/δ_H_ 51.2/4.49), which bear nitrogen (Tables 1 and 2). Overall, **1** was found to possess two rings based on the combinational analysis of the NMR data (Figs. S1–S6) and the degrees of unsaturation inherent in the mass data, implying that this compound belongs to the prenylated isoindolinone class.

**Table 1.** ^13^C NMR (ppm, type) assignments for compounds 1–5.

| Position | 1<sup>a</sup> | 2<sup>b</sup> | 3<sup>b</sup> | 4<sup>b</sup> | 5<sup>c</sup> |
| --- | --- | --- | --- | --- | --- |
| 1 | 167.3 | 191.2 | 191.1 | 171.5 | 192.6 |
| 2 |  | 130.7 | 130.0 |  | 117.4 |
| 3 | 44.5 | 114.2 | 110.1 | 67.6 | 163.5 |
| 3a | 122.4 |  |  | 121.4 |  |
| 4 | 146.7 | 146.5 | 146.6 | 144.9 | 97.0 |
| 5 | 137.6 | 151.3 | 152.1 | 139.0 | 165.5 |
| 6 | 153.8 | 111.4 | 114.5 | 154.3 | 125.6 |
| 7 | 96.5 | 124.6 | 127.6 | 100.0 | 135.3 |
| 7a | 128.2 |  |  | 126.0 |  |
| 8 |  |  |  |  | 16.9 |
| 1' | 68.2 | 66.2 | 66.2 | 69.9 | 25.1 |
| 2' | 120.8 | 118.3 | 118.4 | 119.0 | 122.4 |
| 3' | 139.7 | 143.3 | 143.2 | 144.4 | 143.3 |
| 4' | 38.8 | 39.9 | 39.9 | 39.8 | 40.9 |
| 5' | 25.9 | 26.3 | 26.3 | 26.4 | 27.3 |
| 6' | 123.7 | 123.6 | 123.6 | 123.6 | 124.5 |
| 7' | 134.5 | 135.9 | 135.8 | 135.9 | 136.0 |
| 8' | 38.9 | 39.7 | 39.7 | 39.9 | 40.8 |
| 9' | 26.2 | 26.9 | 26.9 | 26.9 | 27.9 |
| 10' | 124.1 | 124.4 | 124.4 | 124.4 | 125.3 |
| 11' | 130.6 | 131.6 | 131.6 | 131.6 | 132.1 |
| 12' | 25.5 | 25.9 | 25.9 | 25.9 | 26.0 |
| 13' | 17.5 | 17.9 | 17.9 | 17.9 | 17.8 |
| 14' | 15.7 | 16.2 | 16.2 | 16.2 | 16.3 |
| 15' | 16.1 | 17.0 | 16.9 | 16.6 | 16.4 |
| 1'' | 173.8 |  |  |  |  |
| 2'' | 51.2 |  |  |  |  |
| 3'' | 17.2 |  |  |  |  |
| 5-OMe |  |  |  |  | 56.2 |
| 6-OMe | 55.8 |  |  | 56.5 |  |

<sup>a,b,c</sup> Data were obtained in DMSO-*d*<sub>6</sub>, CDCl<sub>3</sub>, and MeOD-*d*<sub>4</sub> respectively.

**Table 2.** ^1^H NMR [δ, mult, (*J* in Hz)] Assignments for Compounds 1–5.

| Position | 1<sup>a</sup> |  |  | 2<sup>b</sup> |  |  | 3<sup>b</sup> |  |  | 4<sup>b</sup> |  |  | 5<sup>c</sup> |  |
| --- | --- | --- | --- | --- | --- | --- | --- | --- | --- | --- | --- | --- | --- | --- |
| 1 |  |  |  | 9.84 | s |  | 9.82 | s |  |  |  |  | 10.36 | s |
| 3 | 4.59 | d | 17.3 | 7.44 | d | 2.0 | 7.43 | s |  | 5.22 | s |  |  |  |
|  | 4.17 | d | 17.3 |  |  |  |  |  |  |  |  |  |  |  |
| 4 |  |  |  |  |  |  |  |  |  |  |  |  | 6.38 | s |
| 6 |  |  |  | 6.97 | d | 8.5 | 7.04 | d | 8.0 |  |  |  |  |  |
| 7 | 6.69 | s |  | 7.41 | dd | 8.0,<br>2.0 | 7.41 | dd | 8.0,<br>1.5 | 7.01 | s |  |  |  |
| 8 |  |  |  |  |  |  |  |  |  |  |  |  | 2.47 | s |
| 1' | 4.46 | d | 6.9 | 4.70 | d | 6.0 | 4.68 | d | 6.5 | 4.71 | d | 7.5 | 3.33 | m |
| 2' | 5.48 | t | 7.0 | 5.49 | t | 7.5 | 5.49 | t | 6.5 | 5.48 | t | 7.0 | 5.03 | m |
| 4' | 1.97 | dt | 8.2,<br>6.5 | 1.98 | m |  | 1.97 | m |  | 1.98 | m |  | 1.88 | m |
| 5' | 2.02 | m |  | 2.14 | m |  | 2.14 | m |  | 2.05 | m |  | 2.07 | m |
| 6' | 5.08 | m |  | 5.08 | dt | 6.5 | 5.10 | dt | 6.5 | 5.08 | m |  | 5.03 | m |
| 8' | 1.93 | dt | 8.3,<br>6.9 | 2.13 | m |  | 2.12 | m |  | 2.06 | m |  | 1.99 | m |
| 9' | 2.00 | m |  | 2.05 | m |  | 2.05 | m |  | 2.05 | m |  | 1.94 | m |
| 10' | 5.06 | m |  | 5.10 | dt | 6.5 | 5.10 | dt | 6.5 | 5.07 | m |  | 4.99 | m |
| 12' | 1.63 | s |  | 1.68 | s |  | 1.68 | s |  | 1.68 | s |  | 1.64 | s |
| 13' | 1.55 | s |  | 1.60 | s |  | 1.60 | s |  | 1.59 | s |  | 1.55 | s |
| 14' | 1.55 | s |  | 1.61 | s |  | 1.61 | s |  | 1.66 | s |  | 1.55 | s |
| 15' | 1.59 | s |  | 1.77 | s |  | 1.76 | s |  | 1.60 | s |  | 1.76 | s |
| 1" |  |  |  |  |  |  |  |  |  |  |  |  |  |  |
| 2" | 4.49 | d | 7.3 |  |  |  |  |  |  |  |  |  |  |  |
| 3" | 1.31 | d | 7.1 |  |  |  |  |  |  |  |  |  |  |  |
| 5-OMe |  |  |  |  |  |  |  |  |  |  |  |  | 3.84 | s |
| 6-OMe | 3.79 | s |  |  |  |  |  |  |  | 3.93 | s |  |  |  |

<sup>a,b,c</sup> Data were obtained in DMSO-*d*<sub>6</sub>, CDCl<sub>3</sub>, and MeOD-*d*<sub>4</sub> respectively.

The COSY and HMBC correlations revealed the characteristic connections within the farnesyl side chain. (Fig. 5). The COSY signals of H_2_-1’–H-2’ (δ_H_ 4.46 and 5.48), H_2_-4’–H-6’ (δ_H_ 1.97, 2.02, and 5.08), and H_2_-8’–H-10’ (δ_H_ 1.93, 2.00, and 5.06) and the HMBC correlations of these protons and four methyl protons at δ_H_ 1.63, 1.55, 1.55, and 1.59 (H_3_-12’–H_3_-15’, respectively) with neighboring carbons revealed the presence of the farnesyl side chain. The HMBC coupling of the methylene protons H_2_-1’ (δ_H_ 4.46) to a quaternary carbon at δ_C_ 137.6 indicated *O*-prenylation of the aromatic system. HMBC couplings of H_2_-3 (δ_H_ 4.59 and 4.17) with C-1 (δ_C_ 167.3), C-3a (δ_C_ 122.4), C-4 (δ_C_ 146.7) and C-7a (δ_C_ 128.2) and H-7 (δ_H_ 6.99) with C-1 (δ_C_ 167.3), C-3a (δ_C_ 122.4), and C-5 (δ_C_ 137.6) revealed the isoindolinone substructure. HMBC coupling of δ_H_ 3.90 and C-6 (δ_C_ 156.0) established the position of the methoxy substituent. The COSY correlations of H-2’’ (δ_H_ 4.49) and H_3_-3’’ (δ_H_ 1.31) and the HMBC correlations of these protons with C -1’’ (δ_C_ 173.8) revealed the alanine moiety. The direct linkage between alanine moiety to the isoindolinone moiety was suggested by a crucial HMBC correlation of α-proton of alanine at H-2’’ (δ_H_ 4.49) and a nitrogenous methylene carbon of isoindolinone at C-3 (δ_C_ 44.5). The planar structure of **1** was found to be similar to aspernidine A, a recently reported prenylated isoindolinone from the fungus *A. nidulans*. Thus, compound **1**, designated nidulene A, was determined to be a new aspernidine-type prenylated isoindolinone attached with an alanine.

**Fig. 5.**
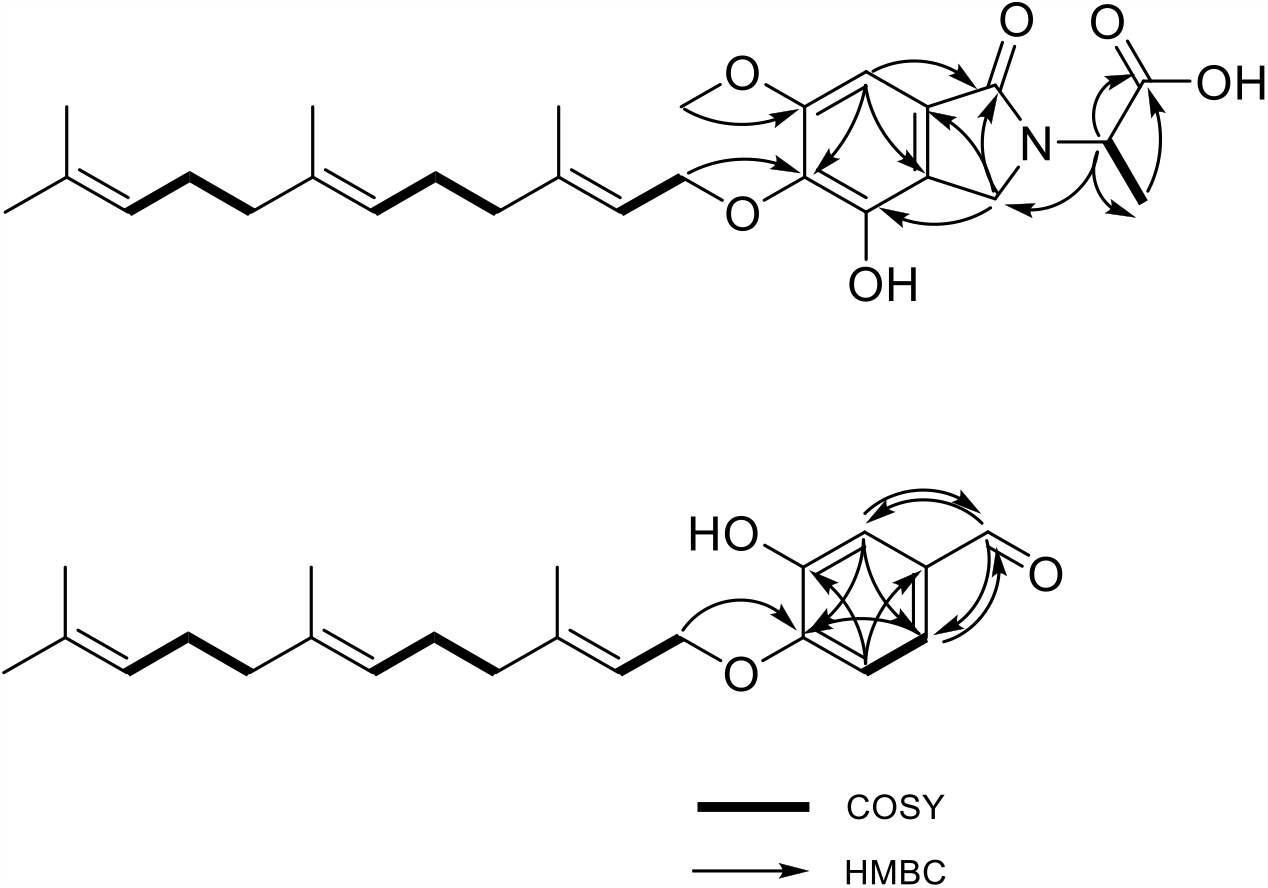
Key correlations of COSY (bold) and HMBC (arrows) experiments for compounds 1–2.

In order to determine the absolute configuration at C-2’’ of **1**, a comparison of the experimental and calculated ECD spectra was carried out. Initially, the calculated ECD results of (*R*)-nidulene A and (*S*)-nidulene A did not show the opposite signs possibly due to the presence of significantly different number of conformers. Total three conformers were obtained from (*R*)-nidulene A while nine conformers were obtained from (*S*)-nidulene A based on a density functional theory (DFT)-based computation. As shown in Fig. 6, the experimental CD data of **1** was almost identical to ECD spectrum of (*R*)-nidulene A. Thus, the absolute configuration at C-2’’ was assigned to *R* configuration and the structural similarity with an amino acid alanine suggest that the amino acid side chain of **1** could be derived from d-alanine.

**Fig. 6.**
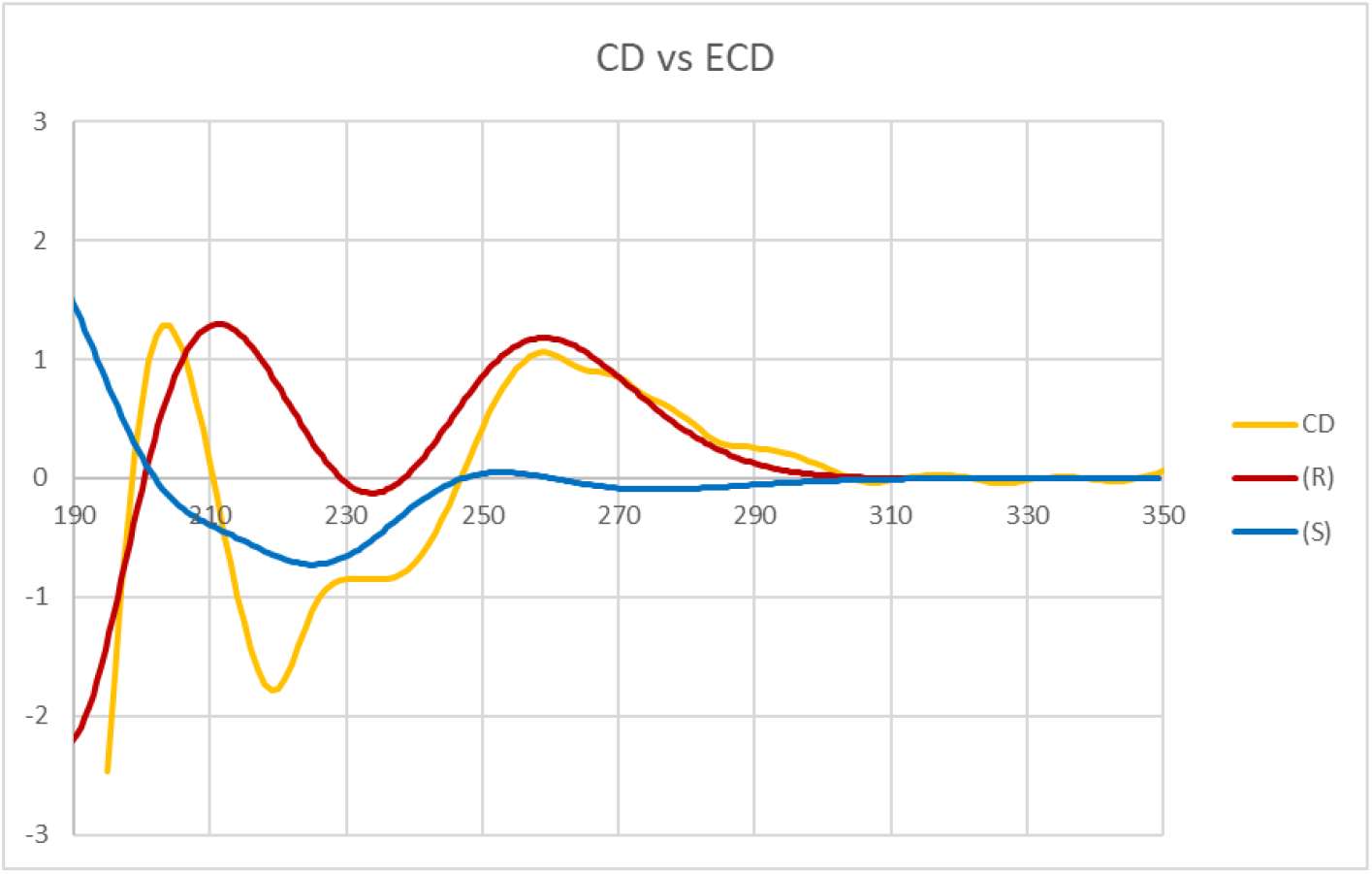
Experimental CD spectrum of nidulene A (1) and calculated ECD spectra of (*R*) and (*S*) forms of nidulene A (1).

Compound **2** was isolated as a yellow amorphous solid with a formula of C_22_H_30_O_3_ based on HR-ESI-MS analysis. The ^13^C NMR data of this compound displayed 22 C-atoms, which were assigned to one aldehyde carbon (δ_C_ 192.2), twelve sp^2^ carbons (δ_C_ 151.3–111.4), four methyl carbons (δ_C_ 25.9–16.2), and five aliphatic carbons (δ_C_ 66.2–26.3) (Table 1, Figs. S7–S11). Same as compound **1**, the COSY and HMBC correlations revealed the farnesyl side chain with 15 C-atoms. Among the remaining seven unassigned carbon atoms, six sp^2^ carbons (δ_C_ 130.7, 114.2, 146.5, 151.3, 114.2, and 124.6; C-2–C-7, respectively) revealed an aromatic ring based on the combination of COSY and HMBC data (Fig. 5). The HMBC correlations from H-1 (δ_H_ 9.84) to C-3 (δ_C_ 114.2) and C-7 (δ_C_ 124.6) showed the evidence of direct attachment at C-2 position of the aromatic ring. The aromatic ring was constructed by the COSY signal between H-6 (δ_H_ 6.97) and H-7 (δ_H_ 7.41) with a coupling constant of 8.0–8.5 Hz and the HMBC correlations from H-6 (δ_H_ 6.97) to C-2 (δ_C_ 130.7) and C-4 (δ_C_ 146.5), H-7 (δ_H_ 7.41) to C-5 (δ_C_ 151.3) and H-3 (δ_H_ 7.44) to C-5 (δ_C_ 151.3) and C-7 (δ_C_ 127.6). The HMBC coupling of the methylene protons H_2_-1’ (δ_H_ 4.70) to a quaternary carbon at C-5 (δ_C_ 151.3) indicated *O*-prenylation of the aromatic ring. Thus, compound **2**, designated nidulene B, was determined to be a new asperugin-type prenylated aromatic derivative.

The ^13^C and ^1^H NMR data (Figs. S12–S13) of compound **3**, which is a known compound (*29*) and has the same molecular formula with **2**, were similar to those obtained for **2**. Both compounds **2** and **3** had the identical carbon and proton chemical shifts for the structure of the farnesyl side chain. The most noticeable difference between **2** and **3** was the replacement of the attachment of a farnesyl chain. For compound **2**, the terpene tail was attached to the *para* position, while compound **3** has a tail at the *meta* position of the aromatic system. Compound **3** was a known compound without proper name (*29*). Thus, compound **3** was named as nidulene C.

The molecular formula of **5** was established as C_24_H_34_O_3_ by HR-ESI-MS analysis. Based on a combination of spectroscopic analyses and a literature survey, compound **5** was identified as a methoxy adduct of the ovinal (Figs S16–S17) (*30*). Although the spectroscopic data of this compound were reminiscent of those of **1**–**4**, several differences were found in the structure of **5**. First, it was found that there is no oxygen bridge between the farnesyl tail and the aromatic head. Also, there was one extra methyl group in the aromatic system than **1**–**4**. Unlike the other four compounds, compound **5** had a direct attachment of methyl to the aromatic head. The planar structure of the compound is the same as ovinal, isolated from *Albatrellus ovinus*, with an extra methoxy group. Thus, the structure of **5**, designated nidulene E, was determined to be a new ovinal derivative (*30*).

In addition to **1**–**3** and **5**, 5 known compounds, prenylated isoindolinone alkaloids (**4, 6, 7**) and linear sesquiterpenes (**8, 9**), were also isolated. Based on a combination of spectroscopic analyses and literature investigations (Figs. S14–S15, S18–S25); these compounds were identified as an aspernidine derivative (**4**) (*29*), aspernidine A (**6**) (*26*), aspernidine F (**7**) (*31*), methyl farnesoate (**8**) (*32*), and JH-diol (**9**) (*33*). The spectroscopic data of these compounds were in good agreement with the reported values. Among these compounds, **4** had not been previously named; therefore, it was named as nidulene D (*29*).

### Human neutrophil chemotaxis is induced by nidulene E (5) and inhibited by nidulene A (1), nidulene C (3), and aspernidine F (7)

As a preliminary examination of crude extracts from TJW336 showed impacts on neutrophil mobility and several studies have identified chemotactic properties of other terpenoids (*34, 35*), we examined all purified metabolites for any effects on neutrophil migration. Upon screening all compounds, nidulene E (**5**) was the only chemoattractive compound. In comparison to the vehicle control (DMSO), we saw increased neutrophil movement to concentrations 0.1, 10, and 100 μg/mL of nidulene E (**5**) (*p* = 0.0134, 0.0081, <0.0001 respectively). Nidulene E (**5**) at 100 μg/mL showed the greatest impact on neutrophil chemotaxis, though less than our positive control fMLP (*p*<0.0001) (Fig. 7A). Inhibition of fMLP-driven neutrophil chemotaxis occurred with the following compounds: nidulene A (**1**) (100 μg/mL, *p =* 0.0009), nidulene C (**3**) (100 μg/mL, *p* = 0.0222), and aspernidine F (**7**) (100 μg/mL, *p* = 0.0001) (Fig. 7B–D). Compounds **2, 4, 8**, and **9** resulted in no chemotaxis or inhibition of chemotaxis across the concentrations tested (Figs. S26 and S27). Aspernidine A (**6**) was tested for both chemotaxis and inhibition but the data was inconclusive due to its high toxicity (data not shown).

**Fig. 7.**
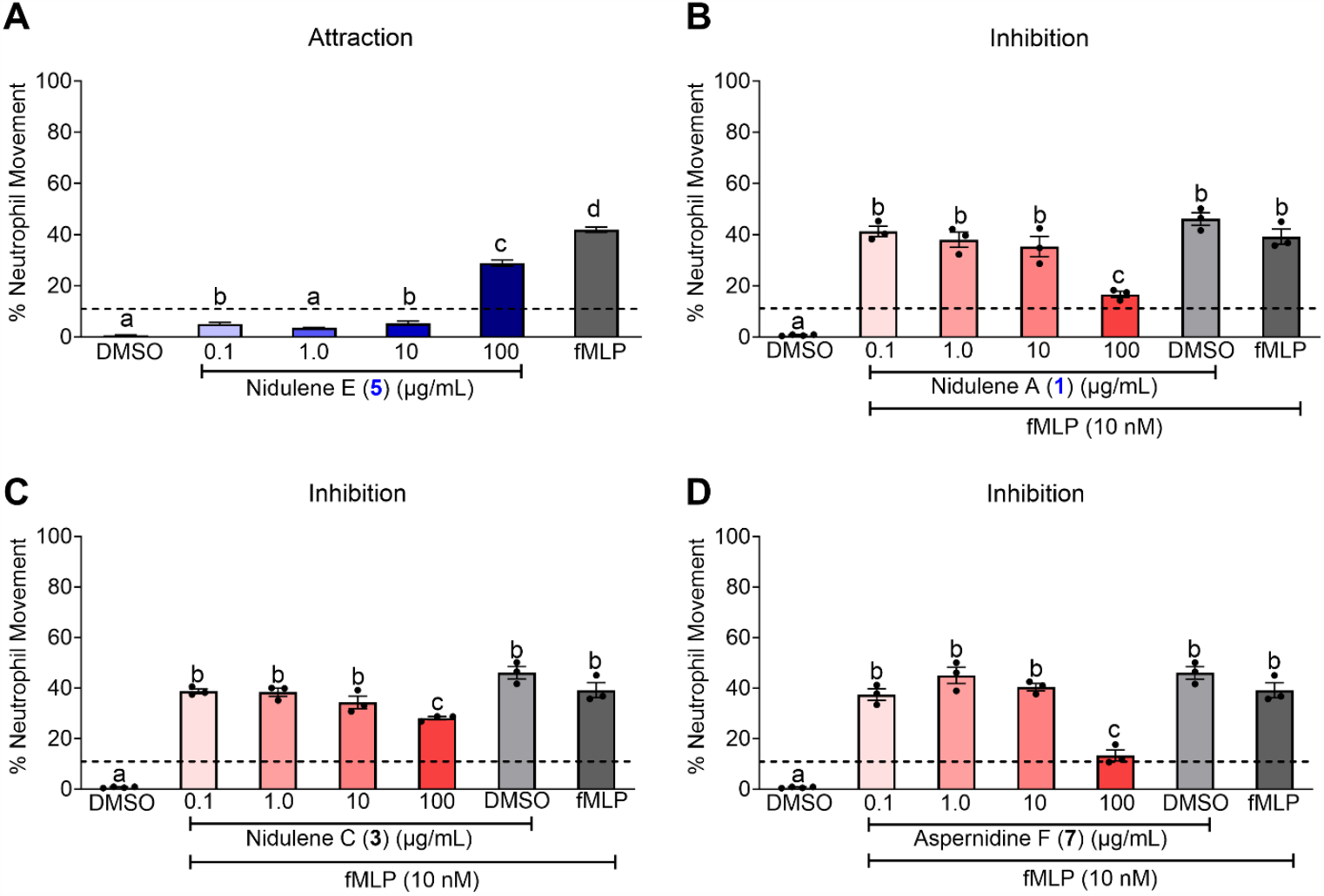
Neutrophil movement is impacted by compounds 1, 3, 5 and 7. **(A)** Nidulene E (**5**) is a chemoattractant for human neutrophils. Using a Boyden chamber, neutrophils stained with calcein AM were seeded in the upper chamber at 5 x 10^4^ cells/well and incubated at 37 °C for 1 h with the vehicle (DMSO; light grey bar), positive control (fMLP, 10 nM; dark grey bar) or the nidulene E (**5**) at four different concentrations (0.1, 1, 10, or 100 μg/mL; blue bars) found in the lower chamber. The media control is indicated as a dotted line across the graph. (**B**–**D**) Inhibition of fMLP-driven neutrophil chemotaxis by nidulene A (**1**), nidulene C (**3**), and aspernidine F (**7**). As described above, neutrophils stained with calcein AM were seeded in the upper chamber at 5 x 10^4^ cells/well and incubated at 37 °C for 1 h with the vehicle (DMSO, light grey bar), the DMSO vehicle with fMLP (fMLP at 10 nM; medium grey bar), positive control (fMLP, 10 nM; dark grey bar) or the respective compounds at four different concentrations (0.1, 1, 10, or 100 μg/mL; red bars) found in the lower chamber. The media control is indicated as a dotted line across the graph. For all graphs (**A**–**D**), cells that migrated into the lower chamber were quantified using flow cytometry and the percent live cells that migrated were determined by calculating the percent live neutrophils from the loading control. Samples are listed as average ± SEM and assays have been repeated to assure reproducibility. Statistical differences were assessed using a one-way ANOVA. Different letters denote statistical differences.

## Discussion

The activation of silent gene clusters in fungi for production of uncharacterized secondary metabolites has garnered significant attention in recent years (*36*). Genetic tools play a crucial role in this pursuit by providing means to unlock the hidden potential of fungal genomes (*37*). One such tool is the implementation of fungal artificial chromosome (FAC) technology that can specifically target and activate silent BGCs in heterologous hosts (*10*). By applying FAC method inversely, we present a twist to FAC utility where we demonstrate that heterologous expression of a *P. destructans* FAC containing a squalene synthase induced cryptic *A. nidulans* aspernidine family compounds including uncharacterized novel immunomodulatory compounds (**1**–**2, 5**; Fig. 1). This activation of aspernidines was accompanied by near loss of austinol, which is considered the primary terpene product in *A. nidulans* (Figs. 2 and 4). The shift of the terpene profile was correlated with shunting of farnesyl pyrophosphate into squalene, thus presenting the potential of terpene manipulation via overexpression of primary metabolite enzymes to redirect flow of the isoprene precursor to different products.

Both farnesyl pyrophosphate and squalene are incorporated into numerous terpenes. Canonical classes include sesquiterpenoids derived from the farnesyl pyrophosphate precursor and triterpenoids derived from squalene (*38, 39*). Logic based on aspernidine structure, therefore, suggested that the linear tail was derived from farnesyl pyrophosphate, but our results suggest the sesquiterpene tail is more likely derived from the triterpene squalene (Fig. 2). Traditionally, terpenes were categorized by the number of isoprenyl units they contained and the origin of terpenes was assumed based on the linear form of isoprenyl units (*40*). However, more recent studies have characterized a variety of terpenes that do not follow the traditional isoprenyl unit rational. For example, triterpenes talaropentaene, macrophomene, and colleterpenol are identified as non-squalene triterpene compounds (*41*). Oxidative degradation of squalene into smaller terpenes has been reported in several publications (*27, 28, 42*). Our work also supports a non-canonical incorporation of squalene into a sesquiterpenoid containing family of metabolites.

The terpenoid potential of *A. nidulans* is only partially defined. Of the ca. 35 terpene synthesis related genes including terpene synthases, terpene cyclases, and/or prenyl transferases, found in the sequenced *A. nidulans* FGSC A4 genome (the isolate used in this study), only nine have been genetically linked to their product: AN1592 and AN1594 (PbcA) to *ent*-pimara-8(14),15-diene, AN9259 (AusN) and AN9257 (AusL) to austinol, AN3228 (PkfE) to aspernidines, AN8514 (tidB) to terrequinone, AN11080 (nptA) to nidulanin A, and AN6784 (xptA) and AN12402 (xptB) to prenylated xanthones (Fig. 2). Products of AN3252, AN6810 and AN9314 are yet to be determined (*43*). Further, there is variation in the genomes of different *A. nidulans* isolates where, for example, isolate SP260548 is missing the austinol clusters (*16*). The results of this study suggest a close relationship between the production of two different *Aspergillus nidulans-*derived sesquiterpenes, austinol and aspernidines, and also implicate the involvement of squalene in the synthesis of aspernidine. By manipulating genes in the host strain, we can gain insights into the correlations of host-derived terpenes and their biosynthetic pathways. Gene manipulation has been instrumental in elucidating terpene discovery in fungi, allowing researchers to gain a deeper understanding of the biosynthetic pathways and regulation of terpene production. For instance, the biosynthetic pathway of the fungal meroterpenoid anditomin, along with its precursors, was clearly identified by heterologous expression of related BGCs from *A. variecolor* in *A. oryzae* (*44*). Moreover, by manipulating potential terpene cyclases in the host strain *Fusarium fujikuroi*, researchers successfully produced and isolated new fungal terpenes, koraiol, and α-acorenol (*45*).

The biological activities of aspernidines are not well characterized with only two studies to-date demonstrating moderate antiproliferative activities of aspernidines A and B (*26*) and anti-tumor properties of aspernidine H in A-594 and SW-480 cell lines (*31*). Based on initial activities of crude extracts of TJW336, we focused on examining any impact of the 9 purified compounds on neutrophil migration. The human neutrophil chemotaxis results supported this initial observation and presented a first look into structure activity relationships of these compounds (Table S2). The aromatic head of these compounds can have immense structural variations, such as isoindolinone, phthalimidine, and phenol and we found the majority of differences in bioactivities attributable to these structural distinctions. Nidulene A (**1**), nidulene C (**3**) and aspernidine F (**7**) inhibited neutrophil chemotaxis whereas nidulene E (**5**) promoted migration. Based on the structural differences matched with the neutrophil migration results, there are several clues that may explain where the activity comes from. For example, Nidulene A (**1**), aspernidine A (**6**), and aspernidine F (**7**) shared the same moiety, which is an isoindolinone but showed different activity. The additional functional groups on the nitrogen from the isoindolinone moiety (compounds **1, 7**) may present inhibition activity toward human neutrophil cells while an NH-proton of the same moiety (compound **6**) may present the cytotoxicity. Unlike other isolated meroterpenes, nidulene E (**5**) showed some chemotaxis activity possibly caused by either the different aromatic head moiety or direct attachment of a linear terpene to the aromatic system without an oxygen bridge. Since the differences from the aromatic head of each terpene showed different activities, we hypothesized that the aromatic head of the terpene may differentiate the inhibition activity on human neutrophil cells.

In conclusion, a tool that shifts the flux of precursors into multiple SM biosynthetic pathways and induce production of novel cryptic compounds has significant potential to drive discovery of novel bioactive compounds. Terpenes encompass a broad class of natural compounds that hold substantial promise for therapeutic applications (*46, 47*). Gaining mastery over the biosynthesis of fungal terpenes opens up new avenues in drug development. Thus, developing methods of controlling and/or altering the production of various terpenes in fungal strains through terpene gene manipulation will add to our understanding of the biosynthetic pathway and biological role of fungal terpenoids (*48*). In this research, we took a different approach using FAC and demonstrated that when a PdFAC1 including *sqsA* was expressed heterologously in *A. nidulans*, it led to changes in the host biosynthesis of endogenous terpene pathways. Compared to the control strain (TJW167), the PdFAC1 transformant (TJW336) exhibited increased production of squalene and compounds belonging to the aspernidine family. We identified three novel and six known fungal terpenes (**1**– **9**). We propose a mechanism in which the conversion of FPP to squalene by *sqsA* leads to increased synthesis of aspernidine type compounds, which may be derived from the squalene at the expense of FPP-derived austinol. We speculate that this method could be employed to successfully modify terpene profiles in other fungi. This research highlights the potential utility of FACs in altering fungal metabolite production and sheds light on the interplay between terpene biosynthetic pathways in filamentous fungi.

## Materials and Methods

### *P. destructans* FAC library

The fungal strains (Table S1) used in this study are *A. nidulans* RJW256 and *P. destructans* 20631-21. *A. nidulans* RJW256 is used as the heterologous host for FAC transformation. For the high molecular weight (HMW) genomic DNA isolation and FAC library construction, *P. destructans* was inoculated from a concentrated spore stock in liquid glucose minimal media (GMM) (*49*). Cultures were incubated at 15 ºC for 2 days and HMW genomic DNA were prepared and FAC library constructed according to the method described previously (*10*). All FAC clones and FAC library are cloned in *Escherichia coli* strain: igMax™ DH10B electrocompetent cells.

### PdFAC1 identification, re-sequencing and key gene deletion via FAC recombineering

Similar to BAC pooling and sequencing (*50*), *P. destructans* FAC library was pooled according to Column_Row_Plate, their pooled DNAs were sequenced by using dual-indexing Illumina sequencing, and DNA assembly, annotation and BGC prediction, and BGC-containing FAC identification with an in-house bioinformatics pipeline. PdFAC1 contained the gene VC83_00068 (genebank#: XM_024463765.1) encoding a putative squalene synthase, SqsA. The *sqsA* was deleted in PdFAC1 using a pair of primers (Table S3) to create the FAC deletant: TJW337 via *E. coli* SW101 using Red/ET tools as previously mentioned (Table S3) (*51, 52*). PdFAC1 was confirmed by re-sequencing and re-annotated.

### Transformation of FACs into the heterologous host strain and construction of TJW336 and TJW337

Transformation in *A. nidulans* was followed as described in a previous study (*53*). Briefly, 2□g of FAC DNA was mixed with protoplasts (10^7^ protoplasts/mL) in presence of 30% PEG-4,000 with 50 mM CaCl_2_ for fungal transformation and transformed protoplast was regenerated on solid GMM with 1.2 M sorbitol and pyridoxine (1 mL of a 0.1% stock solution) as a supplement and incubated it for 3 days at 37 °C in an incubator to obtain transformants. All transformants were confirmed by PCR using a pair of primer listed in Table S3 (data not shown). We constructed the fungal heterologous expression strains: *A. nidulans*/PdFAC1 as TJW336 and *A. nidulans*/PdFAC1Δ*sqsA* as TJW337, respectively.

### Fungal culture and extraction of secondary metabolites

A control strain TJW167, TJW336 and TJW337 were inoculated on triplicated GMM plates and incubated for 7 days at 37 °C according as previously reported (*11*). Subsequently, the entire contents of the plates were collected and lyophilized for 48 h. Samples were then pulverized with a mortar and pestle, and methanol was added. Air-dried methanol extracts were prepared with a SpeedVac system (Savant SpeedVac Concentrator, SC250EXP-115) and then further extracted with organic solvent (chloroform:methanol:ethylacetate, 8:1:1). Organic extracts were evaporated to dryness in a SpeedVac and stored at −20 °C until analysis. For the metabolite isolation and purification, 15 g of crude extract of TJW336 was obtained from 600 plates of solid GMM incubated for 7 days at 37 °C.

### UHPLC–HRMS/MS analysis

Ultra-high pressure liquid chromatography–high resolution mass spectrometry (UHPLC–HRMS) data were acquired using a Thermo Scientific Q Exactive Orbitrap mass spectrometer (Waltham, MA, USA) coupled to a Vanquish UHPLC (Waltham, MA, USA) operated in positive ionization mode. All solvents used were of spectroscopic grade. Extracts from the conditions described below were diluted into 1 mg/mL and used as samples for UHPLC– HRMS/MS. For the general screening including terpenes, a Waters XBridge BEH-C18 column (2.1 × 100 mm, 1.7 μm) was used with acetonitrile (0.05% formic acid) and water (0.05% formic acid) as solvents at a flow rate of 0.2 mL/min. The screening gradient method for the samples is as follow: Starting at 10% organic for 5 min, followed by a linear increase to 90% organic over 20 min, another linear increase to 98% organic for 2 min, holding at 98% organic for 5 min, decreasing back to 10% organic for 3 min, and holding at 10% organic for the final 2 min, for a total of 37 min. For the squalene detection, the same Waters XBridge BEH-C18 column (2.1 × 100 mm, 1.7 μm) was used with methanol and water as solvents at a flow rate of 0.2 mL/min. The screening gradient method for the samples is as follow: Starting at 75% organic followed by a linear increase to 98% organic over 2 min, holding at 98% organic for 15 min, decreasing back to 75% organic for 0.2 min, and holding at 75% organic for the final 2.8 min, for a total of 20 min. A quantity of 10 μL of each sample was injected into the system for the analysis. Each extract was diluted in 1 mg/mL in methanol and 100 ppm of squalene was used as standard.

### General experimental procedures

UV spectra were acquired using a Cary Bio400 UV/Vis spectrophotometer (Varian Inc., Palo Alto, CA, USA). CD spectra were recorded on an AVIV model 420 circular dichroism spectrometer (Hod Hasharon, Israel). IR spectra were recorded on a JASCO 4200 FT-IR spectrometer (Easton, MD, USA) using a ZnSe cell. NMR spectra were recorded in MeOD-*d*_4_, DMSO-*d*_*6*_ or CDCl_3_ solutions on Bruker Avance III HD, 500 MHz instrument (Billerica, MA, USA) equipped with a 5 mm cryoprobe. NMR spectra were processed, and baseline corrected using MestReNova software. UHPLC–HRMS and UHPLC–MS/MS data were acquired using a Thermo Scientific Vanquish UHPLC system (Waltham, MA, USA) connected to a Thermo Scientific Q Exactive Hybrid Quadrupole-Orbitrap mass spectrometer (Waltham, MA, USA) operated in positive and/or negative ionization modes using an *m/z* range of 190 to 2,000. HPLC separations were performed on a Gilson 332 pump and a Gilson 171 DAD detector (Middleton, WI, USA). All solvents used were of spectroscopic grade.

### Compound isolation and purification

The terpene-type compounds from half of the crude extract of TJW336 were initially isolated by preparative reversed-phase HPLC (Waters Xbridge PREP C18 OBD column, 5 μm, 19 × 250 mm) using a gradient solvent system (50% CH_3_CN−H_2_O to 90% CH_3_CN−H_2_O over 35 min, UV detection at 210 nm, and flow rate = 16 mL/min) and afforded compounds **1** (*t*_R_ = 21.8 min), **5** (*t*_R_= 28.4 min), **6** (*t*_R_ = 20.2 min), **7** (*t*_R_ = 17.3 min), **8** (*t*_R_ = 33.3 min), and **9** (*t*_R_ = 17.1 min). Another half of the crude extract was loaded onto the same column using a different gradient solvent system (20% CH_3_CN−H_2_O to 95% CH_3_CN−H_2_O over 24 min, UV detection at 210 nm, and flow rate = 16 mL/min). The fifth fraction (51.2 mg) that eluted with 95% CH_3_CN−H_2_O was separated by preparative reversed-phase HPLC (68% CH_3_CN−H_2_O over 30 min, UV detection at 210 nm, and flow rate = 16 mL/min) and afforded compounds **2** (*t*_R_ = 24.0 min), **3** (*t*_R_ = 26.1 min), and **4** (*t*_R_ = 27.5 min). Compounds **1** was purified by an analytical HPLC (68% CH_3_CN−H_2_O over 30 min; UV detection at 210 nm, and flow rate = 2.0 mL/min; YMC-ODS-A column, 4.6 × 250 mm; *t*_R_ = 22.1 min, respectively). The purified metabolites were isolated in the following amounts: 2.3, 5.6, 5.5, 7.7, 7.6, 3.4, 1.5, 4.4, and 3.3 mg of **1**−**9**, respectively.

*Nidulene A (****1***): yellow, amorphous solid; UV (MeOH) λ_max_ (log ε) 208 (4.24), 263 (1.26) nm; IR (ZnSe) ν_max_ 3247 (br), 1758, 1671 cm^-1^; ^1^H and ^13^C NMR data, Table 1 and 2; HR−ESI−MS *m/z* 472.2684 [M+H]^+^ (calcd for C_27_H_38_NO_6_, 472.2694).

*Nidulene B (****2***): yellow, amorphous solid; UV (MeOH) λ_max_ (log ε) 194 (5.84), 229 (2.34), 276 (1.84), 311 (1.45) nm; IR (ZnSe) ν_max_ 3250 (br), 1734, 1629 cm^-1^; ^1^H and ^13^C NMR data, Table 1 and 2; HR−ESI−MS *m/z* 343.2112 [M+H]^+^ (calcd for C_22_H_31_O_3_, 343.2112).

*Nidulene E (****5***): yellow, amorphous solid; UV (MeOH) λ_max_ (log ε) 205 (5.39), 265 (1.33) nm; IR (ZnSe) ν_max_ 3224 (br), 1729, 1608 cm^-1^; ^1^H and ^13^C NMR data, Table 1 and 2; HR−ESI−MS *m/z* 371.2572 [M+H]^+^ (calcd for C_24_H_35_O_3_, 371.2581).

### ECD calculations

The conformational search for the C-2’’ position of compound **1** was performed using Spartan 14 (v.1.1.7 Wavefunction Inc., Irvine, CA, USA) to identify all possible conformers for each isomer. Through Gaussian 16 software (Wallingford, CT, USA), the equilibrium geometries were optimized to the ground-state level based on the density functional theory (DFT) calculations. The basis parameter set used was B3LYP. Conformers within 15 kJ/mol of each global minimum for *R* and *S* form of **1** were used for gauge-independent atomic orbital (GIAO) shielding constant calculations without geometry optimization at the B3LYP level.

### Human neutrophil chemotaxis to purified compounds (1–9)

Human blood was donated by healthy volunteers based on protocols reviewed and approved by the Institutional Review Board at the University of Wisconsin-Madison. Using a MACSpress neutrophil isolation kit (#130-104-434, MIltenyi) and the erythrocyte depletion kit (#130-098-196, Miltenyi), we isolated human neutrophils from whole blood and stained cells with Calcein AM (C1430, Thermofisher Scientific) at 4 °C for 1h in the dark. Cells were rinsed with PBS and resuspended to 5 × 10^5^ cells/mL. Using a previously established protocol (*54*) with some modifications, a transwell plate (96-well) is prepared 24 h prior to the assay by incubating the upper and lower wells with 10 μg/mL of fibrinogen for 1 h at 37 °C with gentle shaking. Wells are washed with PBS and blocked with 2% bovine serum albumin (BSA) for 30 min at 37 °C before aspiration and drying overnight. On the day of the assay, compounds diluted in mHBSS (1× HBSS, 0.1% human serum albumin, 20 mM HEPES) and added to the respective lower wells at 100 μL/well. Additionally, vehicle controls and fMLP (10 nM) are also prepared in mHBSS and added to requisite wells. Neutrophils are added to the upper chamber at 5 × 10^4^ cells/well and a loading control is used to ensure equal numbers of cells were loaded. Plates are incubated for one hour at 37 °C. Following incubation, we release cells by adding 45 mM EDTA pH 8.0 to lower wells for 15 min. at 4 °C. Samples are transferred to a u-bottom 96-well plate and run on the plate reader of the attune for enumeration following setup of the machine with single-stained samples. The attune is set to mix once prior to drawing up and reading 120 μL of fluid/well. Data is analyzed using FlowJo (10.8.1) software. Percent cell movement is determined by assessing the number of live cells in the treatment well and dividing by the total number of cells recorded in the loading controls.

## Acknowledgments

This work was supported in part by an SBIR award from the National Institute of Allergy and Infectious Diseases at the National Institutes of Health under grant 1R44AI140943 to CCW, JWB and NLK. NPK acknowledges National Institutes of Health R01 [2R01GM112739-05A1 funding SCP and National Institutes of Health R01 AI150669-01A1 funding BNS]. for partial support of this work. We also acknowledge support from National CCIH under grant 5R01AT009143-19 to NPK and NLK. The author(s) thank the University of Wisconsin Carbone Cancer Center Flow Cytometry Laboratory supported by P30-CA014520, for use of its facilities and services. The author(s) also thank the University of Wisconsin National Magnetic Resonance Facility at Madison supported by NIH grant R24GM141526, for use of NMR facilities and the University of Wisconsin–Madison Biophysics Instrumentation Facility supported by the University of Wisconsin–Madison and grants BIR-9512577 (NSF) and S10 RR13790 (NIH) for use of UV and CD facilities.

## Author contributions

Conceptualization: FYL, CCW, JWB, NPK

Methodology: SCP, CCW, JWB

Investigation: SCP, BNS, RG, FAB, HC, RY, TD, JWB

Validation: SCP, BNS, JWB

Formal Analysis: SCP, BNS

Resources: RG, CCW

Supervision: JWB, NPK

Writing—original draft: SCP, JWB, NPK

Writing—review & editing: CCW, NLK, JWB, NPK

## Data and materials availability

All data are available in the main text or the supplementary materials.

